# Blood-based liquid biopsy for comprehensive cancer genomic profiling using next-generation sequencing: an emerging paradigm for noninvasive cancer detection and management in dogs

**DOI:** 10.1101/2021.05.05.442773

**Authors:** Kristina M Kruglyak, Jason Chibuk, Lisa McLennan, Prachi Nakashe, Gilberto E Hernandez, Rita Motalli-Pepio, Donna M Fath, John A Tynan, Lauren E Holtvoigt, Ilya Chorny, Daniel S Grosu, Dana WY Tsui, Andi Flory

**Author notes:** These authors contributed equally to the work and share first authorship (Kristina M Kruglyak, Jason Chibuk). These authors contributed equally to the work and share senior authorship (Dana WY Tsui, Andi Flory).

## Abstract

This proof-of-concept study demonstrates that blood-based liquid biopsy using next generation sequencing of cell-free DNA can noninvasively detect multiple classes of genomic alterations in dogs with cancer, including alterations that originate from spatially separated tumor sites.

Eleven dogs with a variety of confirmed cancer diagnoses (including localized and disseminated disease) who were scheduled for surgical resection, and 5 presumably cancer-free dogs, were enrolled. Blood was collected from each subject, and multiple spatially separated tumor tissue samples were collected during surgery from 9 of the cancer subjects.

All samples were analyzed using an advanced prototype of a novel liquid biopsy test designed to noninvasively interrogate multiple classes of genomic alterations for the detection, characterization, and management of cancer in dogs.

In 5 of the 9 cancer patients with matched tumor and plasma samples, pre-surgical liquid biopsy testing identified genomic alterations, including single nucleotide variants and copy number variants, that matched alterations independently detected in corresponding tumor tissue samples. Importantly, the pre-surgical liquid biopsy test detected alterations observed in spatially separated tissue samples from the same subject, demonstrating the potential of blood-based testing for comprehensive genomic profiling of heterogeneous tumors. Among the 3 patients with post-surgical blood samples, genomic alterations remained detectable in one patient with incomplete tumor resection, suggesting utility for noninvasive detection of minimal residual disease following curative-intent treatment.

Liquid biopsy allows for noninvasive profiling of cancer-associated genomic alterations with a simple blood draw and has potential to overcome the limitations of tissue-based testing posed by tissue-level genomic heterogeneity.

## Introduction

Noninvasive blood-based genomic profiling, often called liquid biopsy, is increasingly employed in human medicine to support cancer diagnosis and management (1,2). It refers to the measurement of circulating biomarkers present in blood that can be used to study the genomic profiles of underlying conditions (e.g., cancer). The most commonly used form of liquid biopsy is based on analysis of cell-free DNA (cfDNA) in plasma; cfDNA is comprised of DNA fragments released by cells into the circulation through secretion, apoptosis or necrosis (3), including cfDNA derived from tumor cells (which is known as circulating tumor DNA or ctDNA). It has been demonstrated in studies of human cancer that plasma cfDNA analysis can recapitulate the genomic profile of the underlying tumor tissue (4,5). Additionally, it offers distinct advantages through its ability to capture the diversity of mutations derived from spatially separated sites within a primary tumor or across metastatic deposits (a fundamental biological feature of cancer, referred to as “tumor heterogeneity”) that may be missed by testing a single tissue sample from a single tumor site (6–8). Liquid biopsy enables genomic profiling of cancer cases that are difficult to biopsy, and its noninvasive nature allows it to be performed safely and repeatedly for disease monitoring. Liquid biopsy has also shown considerable potential as a multi-cancer early detection test that can be used at regular intervals (e.g., annually) for cancer screening in high-risk populations, where detection of the disease at earlier stages can result in improved clinical outcomes (9–12).

A number of studies have demonstrated the presence of ctDNA in canine plasma across multiple cancer types (13–16). The general strategy for detecting tumor-specific cfDNA relies on identifying genomic alterations specific to cancer cells and absent in healthy cells (17). Different cancer types in humans are driven by different classes of genomic alterations (18), a phenomenon that is encountered in canine cancers as well: for instance, single nucleotide variants (e.g., in the *KIT* gene) are frequently found in mast cell tumors (19) whereas copy number variants (gains and losses) are often encountered in canine sarcomas (20).

Here, we report results generated using an advanced prototype of a novel blood-based liquid biopsy assay (hereinafter “the test”) specifically developed for the noninvasive detection, characterization, and management of cancer in dogs. This is the first report of a next-generation sequencing (NGS) based multi-cancer early detection test designed to simultaneously interrogate multiple classes of cancer-associated genomic alterations, including single nucleotide variants (SNVs), insertions and deletions (indels), translocations, and copy number variants (CNVs), in cfDNA extracted from canine blood samples. This proof-of-concept study aims to demonstrate the potential of liquid biopsy to broadly profile tumor-derived genomic alterations, and to capture the genomic heterogeneity of multiple, spatially separated tumor clones (within the same primary site or across different metastatic sites) that would be missed by traditional single-site tissue biopsy, suggesting broad utility as a diagnostic tool in veterinary oncology.

## Methods

Eleven client-owned dogs with a variety of confirmed cancer diagnoses (including localized/regional as well as disseminated/metastatic disease) scheduled for surgical resection, and 5 dogs presumed to be cancer-free (no history of cancer and no clinical signs indicative of cancer at the time of enrollment), were enrolled across multiple care settings. All cancer subjects were enrolled under a dedicated protocol that was independently approved by the institutional animal care and use committees (IACUCs) or internal review boards at two veterinary centers, and all cancer-free subjects were enrolled under a separate dedicated protocol that was independently approved by the institutional clinical review board at one veterinary center; the protocols were also approved by institutional clinical review boards at the other participating sites, based on each site’s requirements. All subjects were client-owned, and written informed consent was obtained from all owners.

From each patient, a baseline blood sample was collected at the time of enrollment. In 9 of the 11 patients with cancer, samples of resected tumor specimens from one or more sites within each primary tumor, and from one or more metastatic sites if available, were collected during the subsequent surgery. In three cancer patients, a post-surgical blood sample was collected 10, 15, and 189 days after surgery, respectively. Clinical and demographic data, plasma volumes, and cfDNA yields for all enrolled subjects are summarized in Table 1.

**Table 1.** Patient demographics and clinical history. FI = female, intact; FS = female, spayed; MI = male, intact; MN = male, neutered; NA = not applicable. ‘Localized/Regional’ for solid tumors designates cancer that is limited to the organ of origin or to nearby lymph nodes, tissues, or organs; ‘Localized/Regional’ for lymphomas designates cancer that is limited to a single lymph node (Stage I) or multiple lymph nodes on one side of the diaphragm (Stage II). ‘Disseminated/Metastatic’ for solid tumors designates cancer that has spread to areas of the body distant from the primary tumor; ‘Disseminated/Metastatic’ for lymphomas designates cancer that involves two or more lymph nodes on both sides of the diaphragm and/or one or more extra-nodal sites (Stages III, IV, and V); ‘Disseminated/Metastatic’ also includes all non-lymphoma hematological malignancies.

| Subject ID | Age (Years) | Sex | Weight (kg) | Reported Breed | Plasma Volume (mL) | Plasma cfDNA Yield (ng) | Primary Diagnosis | Cancer Status | Lymph Node Involvement (Yes/No) | Tumor Size (cm) |
| --- | --- | --- | --- | --- | --- | --- | --- | --- | --- | --- |
| **Cancer patients** |  |  |  |  |  |  |  |  |  |  |
| PT01 | 12 | MI | 9.1 | West Highland White Terrier | 0.6 | 3.1 | Cystic renal carcinoma | Localized/Regional | No | >5 |
| PT02 | 15 | FS | 7.3 | American Eskimo Dog | 8.0 | 6.5 | Cholangiocellular carcinoma | Localized/Regional | No | >5 |
| PT03 | 14 | FS | 18.6 | Mixed | 4.9 | 6.8 | Metastatic pancreatic carcinoma; hepatocellular carcinoma; metastatic splenic hemangiosarcoma | Disseminated / Metastatic | No | >5 |
| PT04 | 11 | FS | 23.2 | Border Collie | 8.0 | 4.1 | Anal sac adenocarcinoma | Localized/Regional | Yes | >5 |
| PT05 | 12 | MN | 27.3 | Mixed | 8.0 | 5.7 | Bilateral anal sac adenocarcinoma | Localized/Regional | Yes | <2 |
| PT06 | 10 | MN | 40.0 | Mixed | 6.6 | 4.2 | Multifocal mast cell tumor | Localized/Regional | No | >5 |
| PT07 | 11 | MN | 36.4 | Labrador | 8.0 | 1.7 | Anal sac adenocarcinoma | Localized/Regional | Yes | <2 |
| PT08 | 9 | FS | 29.1 | Mixed | 8.0 | 2.9 | Soft tissue sarcoma | Localized/Regional | No | 2-5 |
| PT09 | 13 | FS | 23.2 | Mixed | 5.6 | 3.3 | Multifocal soft tissue sarcoma | Localized/Regional | No | >5 |
| PT10 | 8 | MN | 27.7 | Mixed | 9.5 | 1.4 | Osteosarcoma | Localized/Regional | No | 2-5 |
| PT11 | 10 | FS | 24.1 | Goldendoodle | 6.0 | 2.8 | Hemangiosarcoma (renal, splenic) | Disseminated / Metastatic | No | >5 |
| **Presumably cancer-free subjects\*** |  |  |  |  |  |  |  |  |  |  |
| PT12 | 10 | MN | 18.6 | Cardigan Welsh Corgi | 0.5 | 0.4 | NA |  |  |  |
| PT13 | 10 | MN | 7.3 | Miniature Poodle | 5.5 | 2.2 | NA |  |  |  |
| PT14 | 13 | MN | 19.5 | Mixed | 7.5 | 2.1 | NA |  |  |  |
| PT15 | 10 | MI | 53.6 | Mixed | 5.0 | 2.9 | NA |  |  |  |
| PT16 | 13 | FS | 16.8 | Basenji | 6.3 | 2.5 | NA |  |  |  |

\* Subjects presumed to be cancer-free due to no history of cancer and no clinical signs consistent with cancer at the time of blood collection.

Blood samples were processed with a double-centrifugation protocol to separate plasma from white blood cells (WBCs) (21–23). CfDNA was extracted from plasma using a proprietary protocol optimized to maximize cfDNA yield in canine subjects. DNA from WBC and tumor tissue samples was extracted using QIAamp DNA Mini Blood Kit (Qiagen). Amplified DNA libraries were generated for each subject from matched tumor (if available), plasma, and WBC samples using a proprietary method targeting selected regions of the canine genome, followed by next-generation sequencing on an Illumina NovaSeq 6000 for SNV analysis. Amplified libraries were separately subjected to genome-level sequencing for CNV analysis. All sequencing reads were aligned to the CanFam3.1 reference genome (24), and somatic variant calling was performed with a custom bioinformatics pipeline. For subjects who had only a baseline plasma sample available for testing, a proprietary algorithm to detect ctDNA based on fragment length profiles was also applied.

## Results

In the 9 cancer patients that had tumor tissues analyzed, all 9 had somatic alterations detected in tissue, based on aggregate variant calls across all available tissue samples (Supplementary Table 1). In 5 of these subjects, a subset of these variants was also independently detected in the matched pre-operative plasma sample. In 2 cancer patients without a tumor tissue sample available for analysis, both had somatic alterations detected in their respective pre-operative plasma sample, and both were also classified as cancer-positive by fragment length profiling. Among the 7 plasma-positive cancer subjects, 2 had disseminated/metastatic disease and 5 had localized/regional disease (1 with adjacent lymph node involvement and 4 without lymph node involvement). In some patients, only one class of genomic alteration (either SNV or CNV) was detected, suggesting that combining SNV and CNV results improves overall detection of tumor-derived cfDNA compared to relying on just one class of genomic alterations. All 4 remaining cancer subjects who had no variants detected in pre-operative plasma had localized disease (2 with lymph node involvement and 2 without lymph node involvement). These results are consistent with the observation in humans that the proportion of patients with detectable tumor-derived cfDNA in plasma is higher in metastatic disease compared to localized disease (25). No SNV or CNV alterations were detected in plasma samples from any of the 5 presumably cancer-free dogs, and all were also classified as cancer-negative by fragment length profiling. Full subject-level results are summarized in Supplementary Table 1.

The test detected genomic alterations in genes such as *TP53, KRAS, EGFR* and *PIK3CA* that are frequently associated with cancer in both humans and dogs (26,27). Homologs of some of these alterations are clinically actionable in humans; for example, both the plasma and the tissue of a cholangiocarcinoma patient (PT02) featured an *EGFR* p.L805R mutation, which is homologous to the human *EGFR* p.L858R mutation, a common target for EGFR tyrosine kinase inhibitors (28). We also detected mutations in genes that are more specifically associated with canine cancer, such as *PTPN11, PDGFRA* and *PDGFRB;* in particular, the PDGFR protein is a target for the multikinase inhibitor toceranib phosphate (Palladia™) (29,30), which is commonly used in the treatment of canine cancers.

Genomic profiling of spatially separated tumor tissue samples from the same subject demonstrated genomic heterogeneity across sites within the primary mass as well as across distant sites, in keeping with the polyclonal nature of cancer (17); and these different genomic alterations were collectively captured in plasma. For example, in PT04 (diagnosed with anal sac adenocarcinoma with lymph node metastases), tissue samples were collected from two different locations within the left anal sac primary tumor, and two different locations within the left sublumbar lymph node. Four CNVs (on chromosomes 9, 13, 18, 22) are shared across all four samples, while other CNVs are sample-specific: e.g., the CNVs on chromosomes 4 and 27 are specific to the two lymph node samples, while the CNV on chromosome 19 is specific to the two primary tumor samples. The corresponding pre-operative plasma captured most of this heterogeneity, with both shared and private (site-specific) CNV alterations from the various tumor samples reflected in cfDNA (Figure 1).

**Figure 1.**
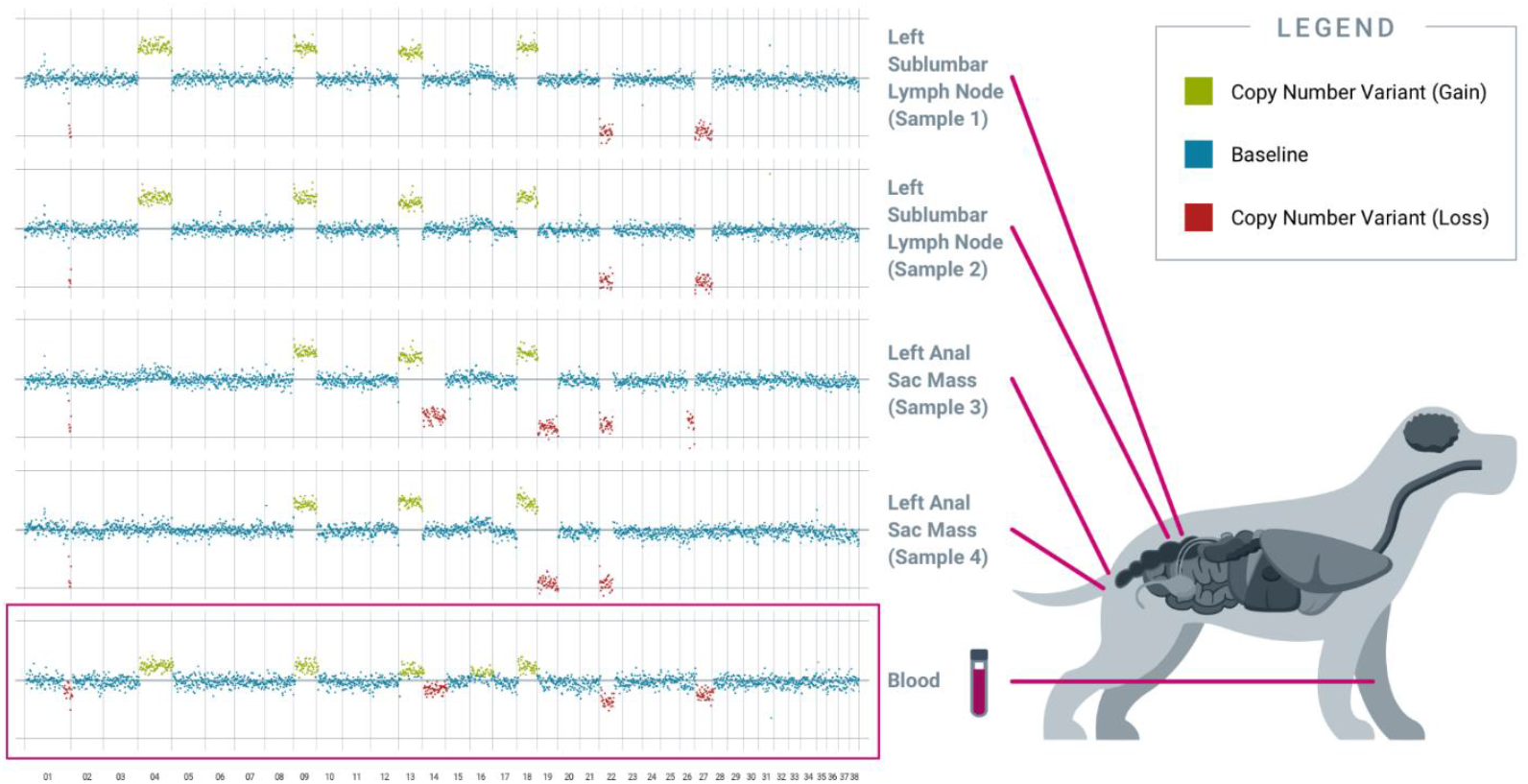
Individual tissue samples reveal distinct somatic copy number alterations, collectively captured in the blood-derived pre-operative plasma. Results from PT04, an 11-year-old female spayed Border Collie diagnosed with apocrine gland anal sac adenocarcinoma of the left anal sac with metastasis to a left sublumbar lymph node, are shown. Four different tumor samples were collected from spatially separated locations at two sites: the left sublumbar lymph node (2 samples) and the left anal sac mass (2 samples). Copy number variation analysis revealed both inter-site and intra-site tumor heterogeneity. Plasma cfDNA captured shared and private alterations from both tumor sites. All chromosomes shown were interrogated for cancer-associated genomic alterations by the test.

Apart from capturing tumor heterogeneity, the test was also able to identify residual disease in plasma after incomplete surgical resection. PT03 was simultaneously diagnosed with multiple primary cancers, including metastatic pancreatic carcinoma, hemangiosarcoma of the liver and spleen, and hepatocellular carcinoma; at surgery, five tissue samples representing primary and metastatic tumor deposits were obtained for testing. Four somatic alterations, including one SNV in *NRAS*, one SNV in *KRAS*, and two distinct SNVs in *TP53*, were identified across four of the five tissue samples (Figure 2), and multiple CNV alterations were also identified in most of the tissue samples. Three of the four SNVs were also detected in the pre-operative plasma sample, along with the majority of the CNV alterations. Complete histologic resection of the hepatic hemangiosarcoma, splenic hemangiosarcoma, and hepatocellular carcinoma were achieved; however, the metastatic pancreatic carcinoma was incompletely excised. This was consistent with the observation of persistent CNV and SNV alterations in the post-operative plasma sample, confirming the presence of residual disease (Figure 3A). For comparison, curative-intent resection of a cystic renal carcinoma in PT01 was complete, and the post-operative plasma sample showed no evidence of any of the somatic alterations that had been present in the pre-operative plasma or the resected tumor (Figure 3B).

**Figure 2.**
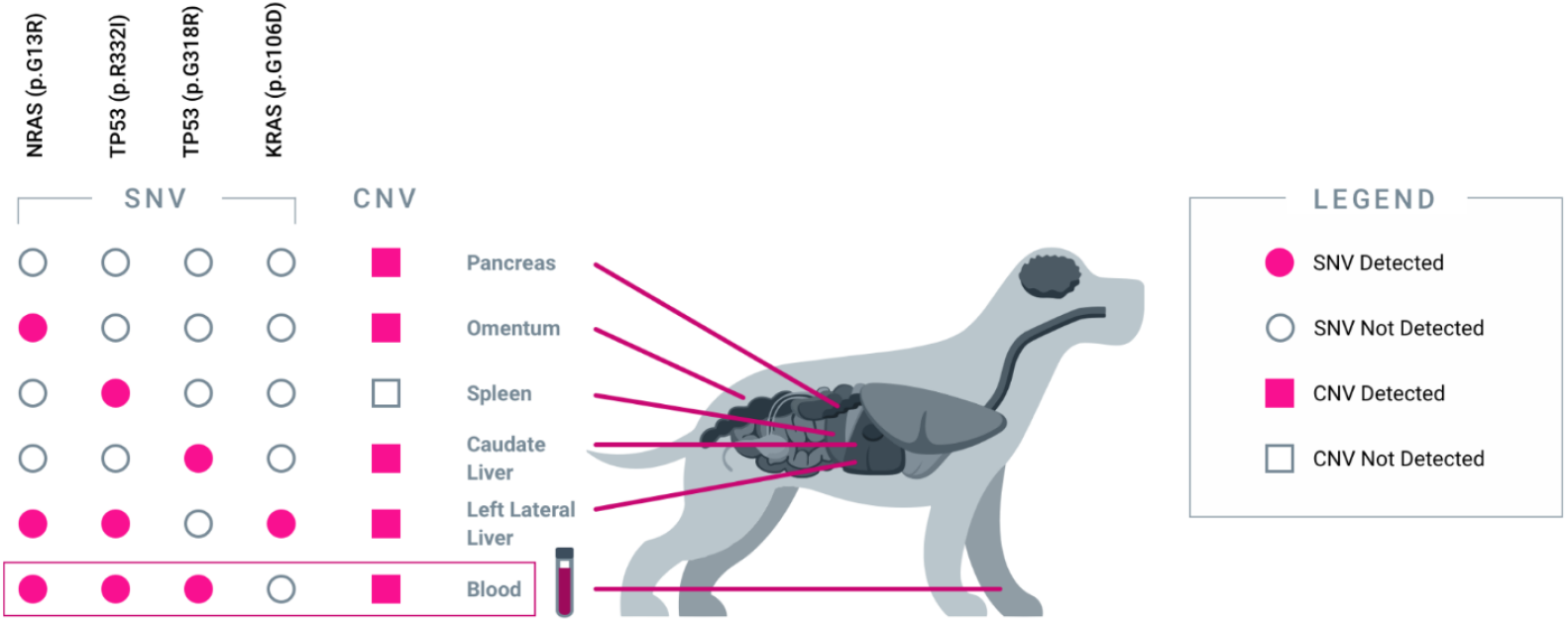
Blood-derived plasma cfDNA captures heterogeneous alterations from five spatially separated sites across multiple organs. Results from PT03, a 14-year-old female spayed mixed breed dog diagnosed with multiple cancers are shown. Five tissue samples were collected from five distinct tumor sites: pancreas (pancreatic carcinoma), omentum (metastatic pancreatic carcinoma), spleen (hemangiosarcoma), caudate liver (metastatic hemangiosarcoma), and left lateral liver (hepatocellular carcinoma). Each tumor sample revealed a unique combination of genomic alterations (SNVs and/or CNVs), some of which were shared across multiple sites. The majority of SNV and CNV alterations were collectively captured in pre-operative plasma cfDNA.

**Figure 3.**
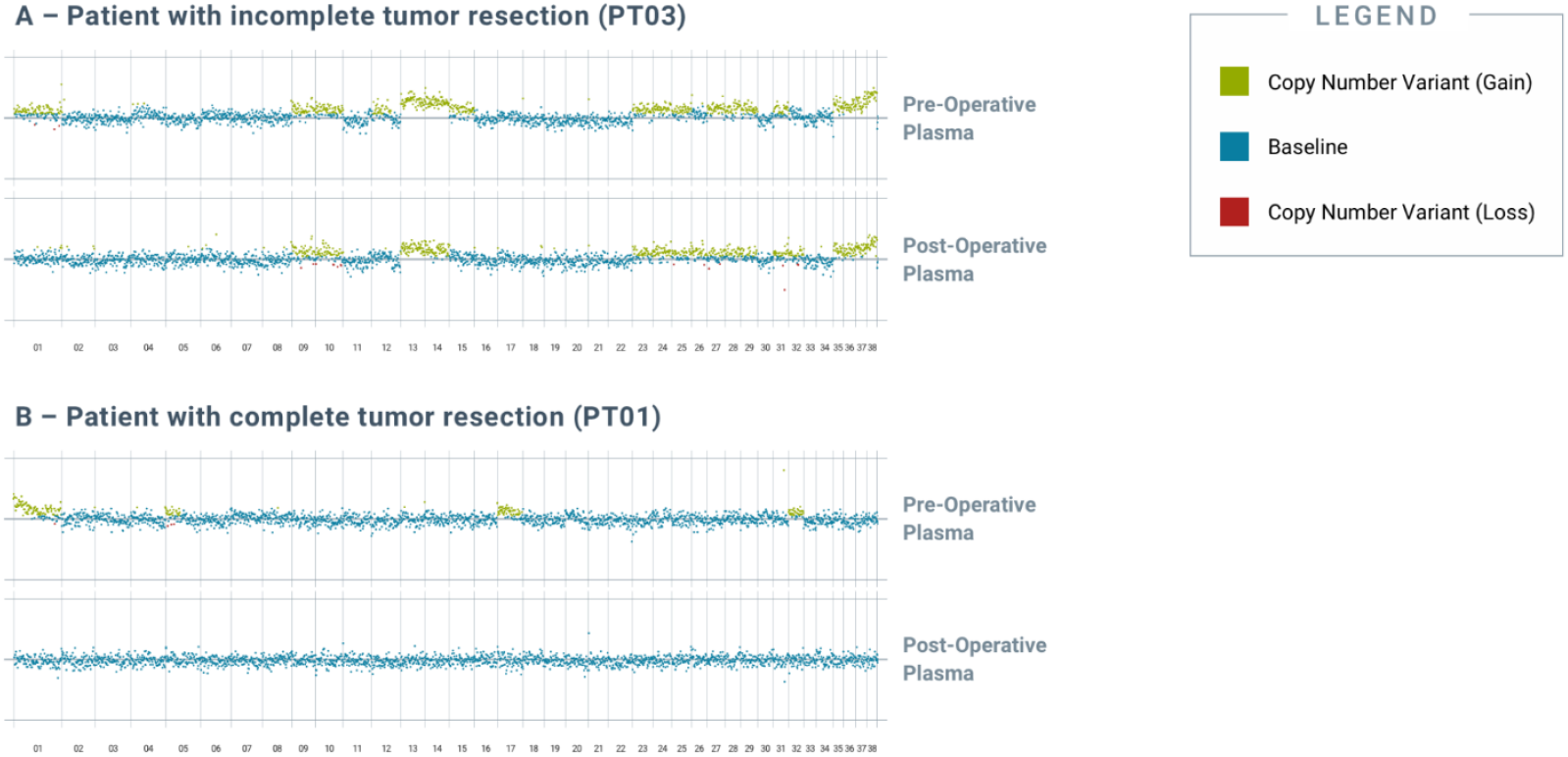
Analysis of post-operative blood-derived plasma cfDNA reveals residual genomic alterations after incomplete resection of tumor. Genomic alterations were detected in post-op blood samples from tumors with incomplete tumor resection but not in those with complete resection. (A) depicts pre- and post-operative plasma CNV traces from PT03, a 14-year-old female spayed mixed breed dog that had incompletely excised metastatic pancreatic carcinoma. (B) depicts pre- and post-operative plasma CNV traces from PT01, a 12-year-old male intact West Highland White Terrier that had completely excised cystic renal carcinoma. All chromosomes shown were interrogated for cancer-associated genomic alterations by the test.

## Discussion

This proof-of-concept study is the first to demonstrate the feasibility of using blood-based liquid biopsy to noninvasively detect multiple classes of genomic alterations in dogs with cancer, in both localized and metastatic settings. An advanced prototype of a novel liquid biopsy test, designed to interrogate multiple classes of cancer-associated genomic alterations across the canine genome, revealed alterations in plasma that were shared between different tumor sites as well as alterations that were private to individual tumor sites within a subject; these observations highlight the potential of plasma cfDNA analysis by NGS to capture the genomic heterogeneity of spatially separated tumors within individual canine cancer patients.

Additionally, our results suggest utility for liquid biopsy in detecting minimal residual disease (MRD) following curative-intent surgery. After a complete resection, tumor-specific genomic alterations should no longer be present in plasma cfDNA; whereas such alterations would persist – potentially at lower levels – following an incomplete resection.

The ability of liquid biopsy to detect genomic alterations originating from one or multiple tumor sites within a patient offers several advantages compared to traditional tissue-based testing. First, it is noninvasive and thus poses a lower risk to patients compared to biopsy or surgery. Second, it allows for a more comprehensive characterization of the full genomic landscape of the patient’s malignancy, across multiple sites within the primary tumor and across metastatic deposits. This tissue-level heterogeneity, which reflects the polyclonal nature of most clinical cancers, would be otherwise difficult to evaluate in dogs due to the morbidity and expense associated with obtaining multiple tissue samples. A blood test to characterize and monitor malignant disease, regardless of its location(s) within the body, is less burdensome and facilitates serial testing for longitudinal monitoring of the disease.

Liquid biopsy testing can support multiple applications across the continuum of canine cancer care, including: screening (for early detection) in high-risk populations, aid in diagnosis, detection of MRD, selection of targeted therapies, and monitoring for cancer recurrence or for treatment response by serial testing; and promises to bring the power of precision oncology to veterinary practice through a simple blood draw that does not require changes to the clinical routine (17). Finally, a liquid biopsy test covering cancer-associated regions of the genome that have high homology between dogs and humans can enable identification of somatic alterations in dogs that have clinically actionable human homologs. Such insights gained from liquid biopsy based genomic profiling could be used to speed the adoption of targeted human cancer therapeutics for the treatment of canine cancer.

Widespread access to multi-cancer early detection testing would be highly beneficial for canine patients, given that cancer is by far the leading cause of death in dogs (31). The current results demonstrate the value of interrogating multiple classes of genomic alterations simultaneously in order to improve sensitivity for detection of cancer. Furthermore, to our knowledge these results are the first to suggest that fragment length profiling may provide utility for cancer detection in dogs. It is important to note that an advanced prototype version of the test was employed in the current study, and further validation in larger cohorts of cancer-diagnosed and cancer-free subjects will be required to fully characterize the analytical performance as well as the clinical sensitivity and specificity of the test.

## Conclusion

Blood-based liquid biopsy testing—first used for noninvasive prenatal testing (32) and more recently for cancer detection and management—is becoming increasingly common in human medicine. Historically, successful advances in human medicine have been quickly adopted for veterinary use after demonstration of utility in companion animals. This study represents the first-ever application of liquid biopsy to simultaneously profile multiple classes of genomic alterations using next-generation sequencing for the noninvasive detection and characterization of cancer in dogs, an important first step toward demonstrating such utility. Additionally, we demonstrate that canine cancers, like human cancers, exhibit significant intra-patient genomic heterogeneity at the tissue level. The sampling challenges imposed by this heterogeneity can be significantly mitigated by a non-invasive liquid biopsy testing approach, which provides a comprehensive, systemic view of the genomic landscape of malignant lesions throughout the body, and which can be easily incorporated into the current clinical routine. Finally, we demonstrate the potential utility of the test for detecting MRD by showing that residual somatic genomic alterations in plasma post-surgery correlate with incomplete surgical resection. Taken together, the findings of this study demonstrate that noninvasive liquid biopsy testing has the potential to support multiple use cases across the continuum of cancer care, for the benefit of canine patients.

### Ethics Statement

The animal study was reviewed and approved by the institutional animal care and use committees (IACUCs) at the University of Guelph Ontario Veterinary College and the University of Minnesota Veterinary Medical Center as well as by institutional clinical review boards at additional sites based on each study site’s requirements; all cancer-free subjects were enrolled under a separate single dedicated protocol, which was independently approved by the Veterinary Centers for America (VCA) Clinical Studies Institutional Review Board as well as by institutional clinical review boards at additional sites based on each study site’s requirements. Written informed consent was obtained from the owners for the participation of their animals in this study.

## Acknowledgements

The authors thank Jason Loftis for assistance with developing the illustrations in this manuscript, Sara Pracht, Sofya Gefter, Daisy Peji, Alma D. Galicia Ortiz, Natasha Shoemaker, Mary Tefend, and Drs. Amber Wolf-Ringwall, Terry Clark, Erin Portillo, K. Scott Griffin, Chelsea Tripp and the doctors and staff at Bridge Animal Referral Centre, and the team at the Mona Campbell Centre for Animal Cancer, OVC, and the Institute for Comparative Cancer Investigation, University of Guelph for sample collections.

