## Supplemental Table 1 for "Blood-based liquid biopsy for comprehensive cancer genomic profiling using next-generation sequencing: an emerging paradigm for noninvasive cancer detection and management in dogs"

Supplementary Table 1. Full subject-level genomic results. Genomic profiling identified somatic alterations in tissue from all cancer positive subjects. Somatic alterations were detectable in the plasma in 7 of 11 cancer -positive subjects. No somatic alterations were detected in the plasma from any presumed healthy subjects.

| Subject Information | | Test Status | | Test Results Detail | | Tumor Biopsy | | | | | Blood Sample | |
|---|---|---|---|---|---|---|---|---|---|---|---|---|
| **Subject ID** | **Primary diagnosis** | **Tumor** | **Blood** | **Detected Variant Type** | **Variant Detail (SNV only)** | **1** | **2** | **3** | **4** | **5** | **Pre Surgery** | **Post Surgery** |
| PT01 | Cystic renal carcinoma | Positive | Positive | SNV | PDGFRA (c.2140+28C>T) | | | | | | NA* | |
| | | | | SNV | ENSCAFG00000047557 (p.Leu117Gln) | | | | | | NA* | |
| | | | | CNV | | | | | | | | |
| PT02 | Cholangiocellular carcinoma | Positive | Positive | SNV | PDGFRB (p.Leu583Pro) | | | | | | | |
| | | | | SNV | EGFR (p.Leu805Arg) | | | | | | | |
| | | | | SNV | CTNNB1 (p.Lys335Ile) | | | | | | | |
| | | | | CNV | | | | | | | | |
| PT03 | Metastatic pancreatic carcinoma; hepatocellular carcinoma; metastatic splenic hemangiosarcoma | Positive | Positive | SNV | NRAS (p.Gly13Arg) | | | | | | | |
| | | | | SNV | TP53 (p.Gly318Arg) | | | | | | | |
| | | | | SNV | KRAS (p.Gly106Asp) | | | | | | | |
| | | | | SNV | TP53 (p.Arg332Ile) | | | | | | | |
| | | | | SNV | PTPN11 (p.Gly507Val) | | | | | | | |
| | | | | CNV | | | | | | | | |
| PT04 | Anal sac adenocarcinoma | Positive | Positive | CNV | | | | | | | | |
| PT05 | Bilateral anal sac adenocarcinoma | Positive | Negative | CNV | | | | | | | | |
| PT06 | Multifocal mast cell tumor | Positive | Negative | CNV | | | | | | | | |
| PT07 | Anal sac adenocarcinoma | Positive | Negative | SNV | PIK3CA (p.Tyr1021Cys) | | | | | | | |
| | | | | CNV | | | | | | | | |
| PT08 | Soft tissue sarcoma | Positive | Positive | CNV | | | | | | | | |
| PT09 | Multifocal soft tissue sarcoma | Positive | Negative | CNV | | | | | | | | |
| PT10 | Osteosarcoma | NA | Positive | SNV | CALR (c.1054-37G>A) | | | | | | | |
| | | | | SNV | TP53 (p.Glu71SerfsTer7) | | | | | | | |
| | | | | CNV | | | | | | | | |
| | | | | Fragment Length | | | | | | | | |
| PT11 | Hemangiosarcoma (renal, splenic) | NA | Positive | SNV | PIK3CA (p.Asn347Lys) | | | | | | | |
| | | | | Fragment Length | | | | | | | | |
| PT12** | Presumed cancer-free | NA | Negative | None | | | | | | | | |
| PT13** | Presumed cancer-free | NA | Negative | None | | | | | | | | |
| PT14** | Presumed cancer-free | NA | Negative | None | | | | | | | | |
| PT15** | Presumed cancer-free | NA | Negative | None | | | | | | | | |
| PT16** | Presumed cancer-free | NA | Negative | None | | | | | | | | |

* The baseline blood sample for PT01 failed minimum coverage requirements.

**Subjects PT12-PT16 were presumed to be cancer-free due to no history of cancer and no clinical signs consistent with cancer at the time of blood collection.

CNV = copy number variant; SNV = single nucleotide variant; NA = not applicable

**LEGEND**

- Variant detected
- Sample at given timepoint not available
- No variant detected
