## Supplemental Figure 1 for "Blood-based liquid biopsy for comprehensive cancer genomic profiling using next-generation sequencing: an emerging paradigm for noninvasive cancer detection and management in dogs"

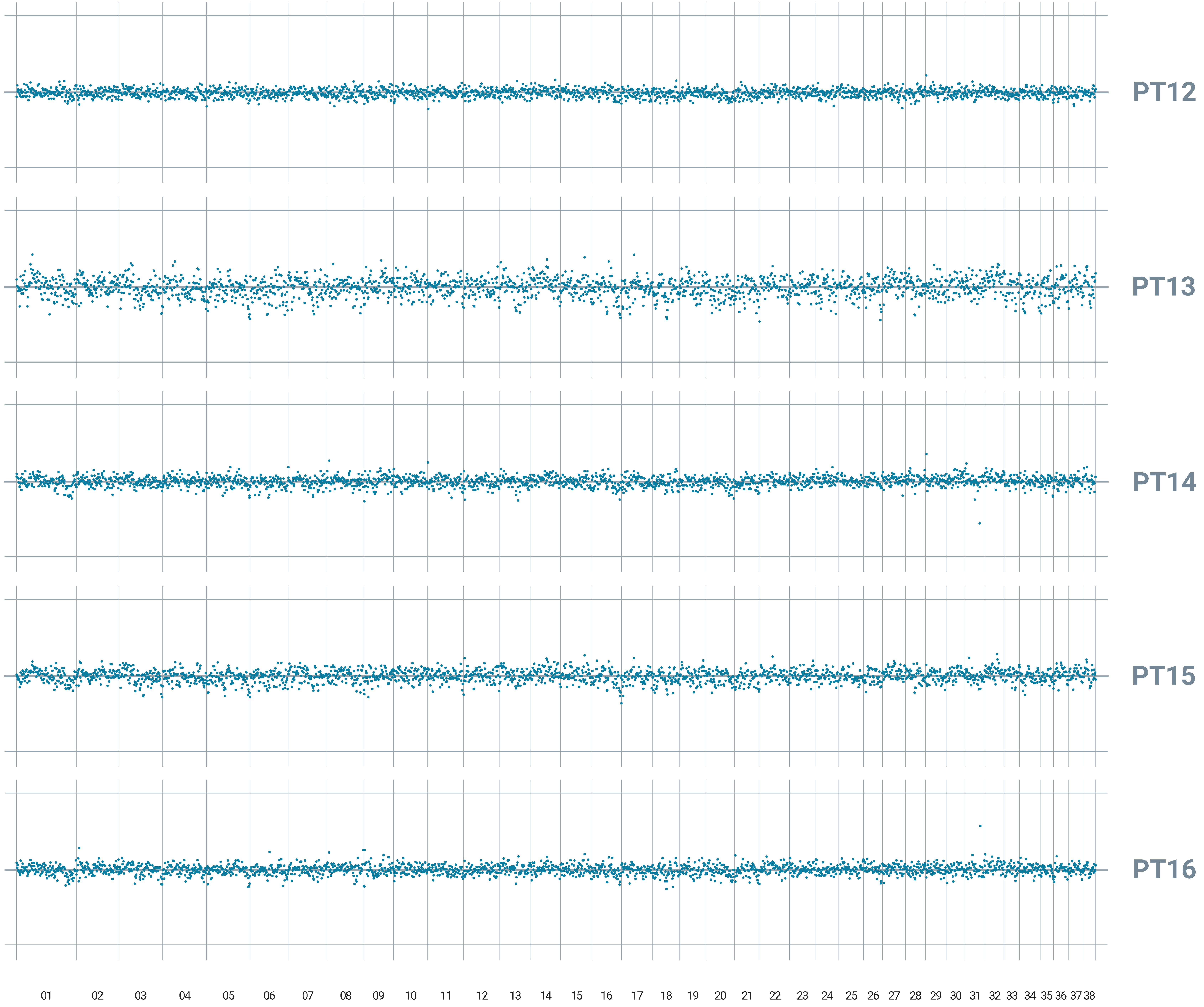


### LEGEND

- Copy Number Variant (Gain)
- Baseline
- Copy Number Variant (Loss)

#### SUPPLEMENTARY FIGURE 1

*Absence of somatic genomic alterations in plasma collected from presumably cancer-free subjects*. No CNV or SNV alterations were detected in the five presumably cancer-free subjects. All chromosomes shown were interrogated for cancer-associated genomic alterations by the test.
